# HyARQ: a Snakemake workflow for allele-specific read counting and quality control in F1 hybrids

**DOI:** 10.64898/2026.10.16.759667

**Authors:** Sontosh K. Deb, Sarah Siegle, Joseph L. Gage

## Abstract

**Summary:** Allele-specific analysis in F1 hybrids allows direct assessment of how cis-regulatory variation contributes to differences in gene expression and other regulatory features. However, allele-specific analyses are sensitive to errors arising from multiple sources, including mapping bias and technical variation, requiring careful processing and quality control. These challenges are compounded by the fact that workflows for different molecular assays are often developed separately, making consistent analysis and quality control across assays difficult. Here, we introduce HyARQ, a Snakemake workflow for allele-specific read counting and quality control across RNA-seq, MOA-seq, ATAC-seq, and ChIP-seq data. HyARQ integrates assay-specific preprocessing, mapping to concatenated parental genomes, filtering, counting at paired parental genes or regulatory regions, and quality control within a common workflow. Using simulated datasets with known parental origin and allelic proportions, we show that HyARQ accurately recovers expected parental proportions with minimal cross-parent assignment across all four assay types. We further applied HyARQ to real RNA-seq, MOA-seq, ATAC-seq, and ChIP-seq datasets and generated paired parental counts and assay-specific quality control outputs for each assay. HyARQ provides a standardized and reproducible framework for allele-specific analysis across multiple molecular layers and facilitates downstream studies of cis-regulatory variation in F1 hybrids.

**Availability and implementation:** HyARQ is available as open-source software, with the source code and example test data hosted at https://github.com/gagelab/hyarq. Permanent archived versions of the HyARQ v1.0.0 software and associated validation data are available through Zenodo at https://doi.org/10.5281/zenodo.23349373 and https://doi.org/10.5281/zenodo.23372517, respectively.

**Supplementary information:** Supplementary data are available with this preprint.

## 1 Introduction

Gene expression is a complex molecular trait that varies extensively among individuals in natural populations (Pastinen 2010, Shao *et al*. 2019, Tsouris *et al*. 2024). Differences in gene expression can arise from cis-regulatory variation, such as changes in promoter or enhancer sequences; from trans-acting factors, such as transcription factors or other diffusible regulators; or from a combination of both (Pastinen 2010, Rojas and Lang 2026, Zhao *et al*. 2026). Allele-specific analysis in F1 hybrids derived from inbred parents provides a powerful approach to dissect the regulatory basis of gene expression (Rojas and Lang 2026). In an F1 hybrid, the two parental alleles share the same cellular environment and are exposed to the same trans-acting factors, so differences in their expression are generally attributed to cis-regulatory divergence (Zhao *et al*. 2026).

Differences in expression between parental alleles, commonly referred to as allele-specific expression (ASE), have been widely studied in F1 hybrids to examine cis-regulatory variation across diverse organisms (Ballinger *et al*. 2023, Hartwig *et al*. 2023, Puixeu *et al*. 2023, Tsouris *et al*. 2023, Engelhorn *et al*. 2025, Deb *et al*. 2026, Huang *et al*. 2026). Regulatory variation in non-coding regions can alter chromatin accessibility, transcription factor binding, or other regulatory processes that ultimately contribute to ASE (Pastinen 2010, Cleary and Seoighe 2021). However, ASE provides a downstream readout of cis-regulatory divergence and does not by itself reveal the molecular mechanisms underlying these differences. Allele-specific measurements of additional regulatory layers can help connect regulatory variation with changes in gene expression and downstream phenotypes.

Recent studies have examined allele-specific chromatin accessibility, transcription factor binding and occupancy, and other molecular phenotypes alongside gene expression, providing a more complete view of cis-regulatory variation (Hartwig *et al*. 2023, Engelhorn *et al*. 2025, Minow *et al*. 2025, Huang *et al*. 2026). As allele-specific analyses expand across these regulatory layers, applying consistent approaches for allele-specific quantification across different molecular assays becomes increasingly important (Mattevi *et al*. 2025).

Several computational tools have been developed for allele-specific analysis, including WASP (Geijn van de *et al*. 2015), SNPsplit (Krueger and Andrews 2016), MEA (Albert *et al*. 2018), ASET (Wu *et al*. 2025), and MIXALIME (Buyan *et al*. 2025). Although these tools differ in scope, many rely on known genetic variants for allele assignment, correction of mapping bias, or construction of personalized or pseudogenome references. An alternative approach is to map reads directly to a concatenated reference containing both parental genome assemblies. This strategy has been used independently to quantify haplotype-specific transcription factor binding with ChIP-seq, transcription factor occupancy with MOA-seq, gene expression with RNA-seq, and chromatin accessibility with ATAC-seq in plant hybrids (Hartwig *et al*. 2023, Engelhorn *et al*. 2025, Minow *et al*. 2025).

Comparison of mapping strategies has shown that a dual parent reference can produce balanced haplotype mapping and substantially reduce parental reference bias compared with single reference and SNP replaced approaches (Engelhorn *et al*. 2025). However, these applications have largely relied on study- and assay-specific analysis pipelines. When high-quality parental genome assemblies are available, concatenated reference mapping provides a practical approach for quantifying allele-specific signals directly from both parental genomes. A standardized framework that applies this strategy to corresponding genes and regulatory regions across multiple molecular assays would facilitate consistent analysis of cis-regulatory variation in F1 hybrids.

Here, we present HyARQ, a Snakemake workflow for quantification and quality control of allele-specific molecular traits in F1 hybrids. HyARQ maps sequencing reads to a concatenated parental reference and quantifies reads over corresponding features from the two parental genomes, allowing differences between parental alleles to be measured at the gene or regulatory region level. The workflow supports RNA-seq, MOA-seq, ATAC-seq, and ChIP-seq, with assay-specific processing followed by a common framework for allele-specific quantification and reporting. HyARQ generates paired feature level parental counts, assay wide count matrices, and quality control reports for each assay. By standardizing quantification and reporting across multiple molecular data types, HyARQ provides a consistent framework for examining cis-regulatory differences across regulatory layers.

## 2 Materials and Methods

### 2.1 Workflow overview and input data

The HyARQ workflow consists of five steps: 1) dual parent reference preparation and paired feature validation, 2) read preprocessing, 3) assay-specific alignment and filtering, 4) feature level counting, and 5) QC reporting (**Figure 1**). Required inputs include paired end raw FASTQ files, two parental reference genomes, gene annotations (RNA-seq only), and assay-specific paired feature files linking each gene or peak region in one parental genome to its allelic counterpart in the other. Workflow settings are provided through (i) a YAML file containing the parental resources and prefixes, assay-specific options, filtering thresholds, computational resources, and output settings, and (ii) a tab-delimited sample metadata file linking library IDs to assay type, replicate information, and paired FASTQ files. This metadata file determines which assay modules and libraries are processed

**Figure 1.**
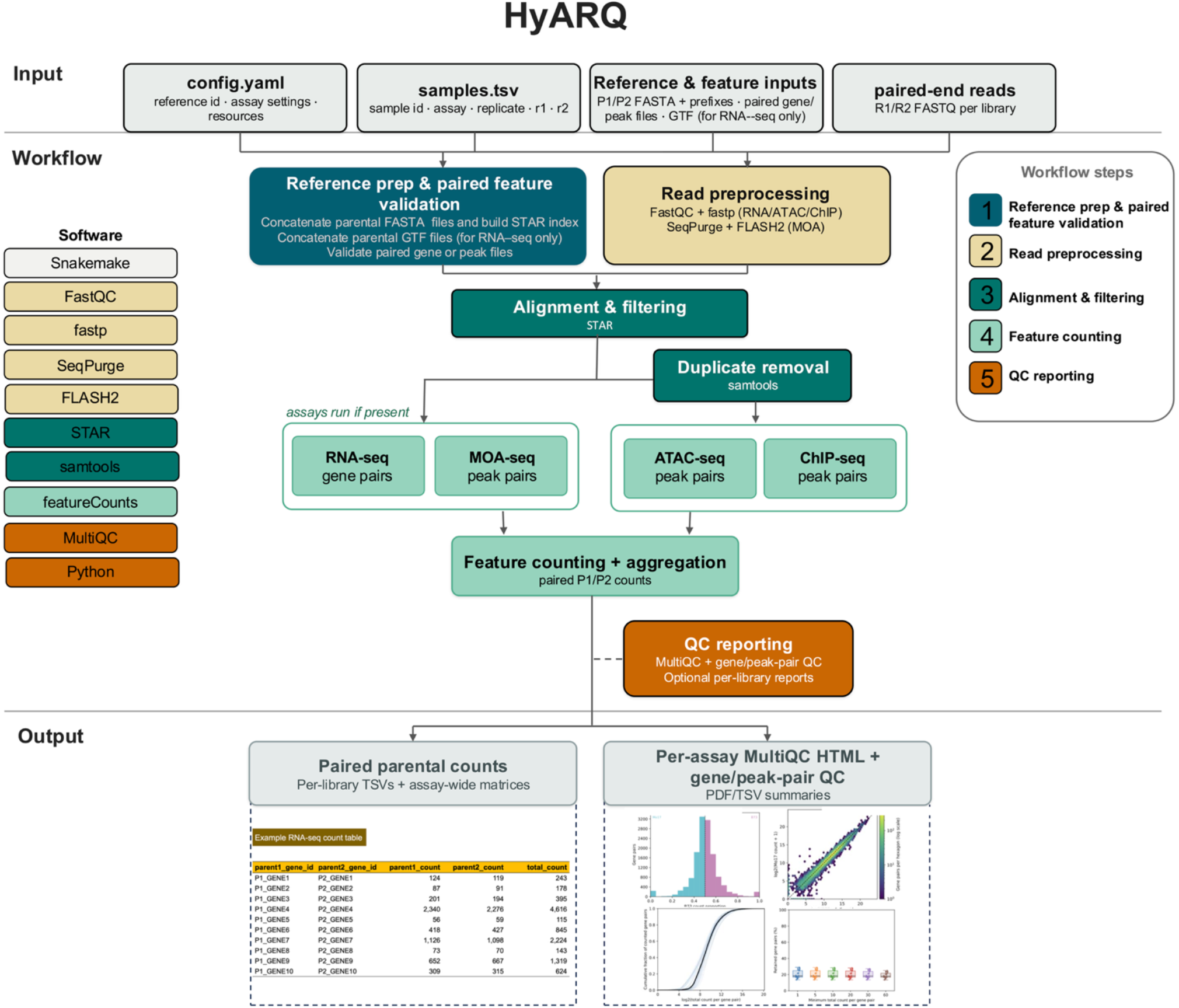
Overview of the different steps in the HyARQ workflow. HyARQ takes paired-end FASTQ files, two parental reference genomes, assay-specific paired feature files, and gene annotations for RNA-seq as input. Workflow settings and sample information are provided through config.yaml and samples.tsv. The workflow consists of five main steps: reference preparation and paired feature validation, read preprocessing, alignment and filtering, feature counting, and QC reporting. Parental FASTA files are concatenated to build the STAR index, and parental GTF files are concatenated when RNA-seq is analyzed. RNA-seq, ATAC-seq, and ChIP-seq reads are preprocessed with FastQC and fastp. MOA-seq paired-end reads are processed with SeqPurge and merged into single-end reads with FLASH2 before alignment. RNA-seq reads are aligned with STAR using splice-aware two-pass mapping, whereas MOA-seq, ATAC-seq, and ChIP-seq are aligned with spliced alignment disabled (--alignIntronMax 1). Duplicate removal is applied to ATAC-seq and ChIP-seq. Assay modules are run only when present in the sample metadata. RNA-seq is quantified at paired genes, while MOA-seq, ATAC-seq, and ChIP-seq are quantified at paired peak regions. HyARQ outputs per-library parental count tables, assay-wide count matrices, MultiQC reports, and gene- or peak-pair QC summaries; optional per-library QC reports can also be generated. The lower panels show examples of a parental count table and assay-level QC output.

### 2.2 Dual parent reference preparation and paired feature validation

Chromosome and contig names in the two parental reference genomes are prefixed with user provided Parent1 and Parent2 identifiers (e.g., for a cross between parents B73 and Mo17, B73_chr1 and Mo17_chr1) before the genomes are concatenated. For RNA-seq, users also provide gene annotations in GTF format, which are updated with the same parental prefixes and concatenated. HyARQ validates the dual parent reference and annotation files by checking that sequence names are unique, prefixes are applied consistently, annotation sequence names match the corresponding FASTA records, and annotation records do not refer to sequences missing from the reference. HyARQ also checks that the user-provided paired feature file has the required structure and is suitable for downstream counting.

### 2.3 Read preprocessing

Raw paired-end FASTQ reads are assessed with FastQC (Andrews 2010). RNA-seq, ATAC-seq, and ChIP-seq reads are trimmed and filtered with fastp (Chen *et al*. 2018). MOA-seq reads are processed with SeqPurge (Sturm *et al*. 2016), and overlapping read pairs are then merged with FLASH2 (Magoč and Salzberg 2011) into single-end reads before alignment. Processed reads from all assay types are reassessed with FastQC. Tool parameters are defined separately for each assay.

### 2.4 Assay-specific alignment and filtering

RNA-seq reads are aligned to the concatenated parental reference with STAR (Dobin *et al*. 2013) using splice-aware two-pass mapping. For ATAC-seq, ChIP-seq, and MOA-seq, --alignIntronMax 1 is used to prevent spliced alignments. ATAC-seq and ChIP-seq reads are mapped in paired-end mode, whereas merged MOA-seq reads are mapped as single-end. Uniquely mapped alignments are retained for all assays, with additional assay-specific filtering (additional information in Supplementary Note, section 2.4).

### 2.5 Assay-specific parental assignment and feature level counting

HyARQ uses featureCounts (Liao *et al*. 2014) to generate parental read counts at predefined paired features (additional information in Supplementary Note, section 2.5). For RNA-seq, gene counts are generated using the concatenated parental GTF annotation and matched according to the supplied gene pair file. For MOA-seq, ATAC-seq, and ChIP-seq, counts are generated at supplied paired peak regions in the two parental genomes. MOA-seq is counted from merged single-end alignments, whereas ATAC-seq and ChIP-seq are counted as paired-end fragments. For each gene or peak pair, HyARQ reports the Parent1 count, Parent2 count, and total count and combines counts across libraries into assay-level matrices.

### 2.6 QC reporting

For each assay type, MultiQC (Ewels *et al*. 2016) aggregates preprocessing and alignment matrices into an HTML report summarizing read quality, filtering, and mapping. This report can be used to identify libraries with poor read quality, excessive read loss, or low mapping performance. HyARQ also generates assay-specific QC figures from the corresponding paired feature count tables. These figures summarize the number of paired features with detectable counts, count depth, parental count proportions, and feature retention at selected count thresholds for individual samples and for the experiment as a whole. The underlying summary statistics are also provided as tab-delimited files. The assay-specific QC figures and summary tables can be used to compare feature recovery, count depth, and parental balance among samples and to evaluate how different minimum count thresholds affect the number of gene or peak pairs retained for downstream analyses.

## 3 Results

To evaluate HyARQ across the four supported assay types, we used 1) simulated datasets with known parental origin and allelic proportions, and 2) real RNA-seq, MOA-seq, ATAC-seq, and ChIP-seq datasets. The simulations were used to evaluate parental assignment accuracy and fragment recovery, whereas the real datasets were used to assess the assay-specific workflows and generation of paired parental counts. Detailed descriptions of the datasets and quality control summaries are provided in Supplementary Note.

### 3.1 Testing with simulated data

To test HyARQ with known parental origin and allelic proportions, we generated synthetic RNA-seq, MOA-seq, ATAC-seq, and ChIP-seq datasets from two parental genomes containing corresponding loci with sequence differences. Four paired features were simulated for each assay with expected Parent1 fractions of 0.50, 0.80, 0.20, and 0.50. RNA-seq features consisted of 1.5-kb genes with two exons, whereas ATAC-seq, MOA-seq, and ChIP-seq features were represented by 500-bp, 80-bp, and 400-bp paired regions, respectively. Two replicates were generated for each assay and processed through the complete HyARQ workflow.

HyARQ accurately recovered the simulated parental fractions across all four assays. The mean absolute error between expected and observed Parent1 fractions was 0.0020 for RNA-seq, 0.0047 for MOA-seq, 0.0017 for ATAC-seq, and 0.0051 for ChIP-seq, with a maximum error of 0.0231 across all feature-replicate combinations. Cross-parent assignment was low, ranging from 0% for RNA-seq to 0.11% for MOA-seq. Mean final fragment recovery was 96.6% for RNA-seq, 77.4% for MOA-seq, 93.8% for ATAC-seq, and 84.9% for ChIP-seq. Although recovery varied among assays and features, the expected parental fractions were maintained.

### 3.2 Testing with real data

For RNA-seq, we applied HyARQ to six B73×Mo17 hybrid samples (Deb *et al*. 2026). A total of 24,611 paired parental genes were used as input, and 15,624 gene pairs had counts in at least one sample. The overall Parent1 fraction was 0.493, indicating no strong overall bias toward either parental genome.

For MOA-seq and ChIP-seq, we applied HyARQ to three replicates of publicly available B73×Mo17 hybrid datasets from Engelhorn et al. (2025) and Harwig et al. (2023), respectively. The ChIP-seq dataset measured binding of the brassinosteroid-responsive transcription factor ZmBZR1 using a hybrid allele-specific chromatin-binding approach. From the peak sets reported by these studies, we generated one-to-one paired parental peak regions for use as counting units. Counts were obtained for all 8,242 MOA-seq paired peak regions, with an overall Parent1 fraction of 0.500. Similarly, counts were obtained for all 1,927 paired ZmBZR1 ChIP-seq peak regions, with an overall Parent1 fraction of 0.493.

For ATAC-seq, due to lack of publicly available plant hybrid data, we used B73 and CML103 inbred sequencing data and accessible chromatin regions from Hufford et al. (2021). We generated four synthetic B73×CML103 hybrid samples by randomly sampling equal numbers of paired-end reads from the two parental datasets and generated 14,302 one-to-one paired accessible chromatin regions from the parental peak sets. Of these, 12,300 paired regions had counts in at least one synthetic hybrid sample, with an overall Parent1 fraction of 0.530. The parental proportions estimated from paired region counts closely followed the genome-wide parental proportions observed after mapping across the four synthetic hybrid samples, consistent with library-specific differences in source coverage and read distribution rather than a systematic parental bias introduced by HyARQ.

Workflow runs were performed on a high-performance computing cluster using Slurm. Complete HyARQ execution required 2.07 h (34.4 CPU-h) for RNA-seq, 0.77 h (13.8 CPU-h) for MOA-seq, 0.96 h (8.9 CPU-h) for ATAC-seq, and 0.90 h (6.0 CPU-h) for ChIP-seq. Peak memory used by any individual job was 51.4 GB for RNA-seq, 45.3 GB for MOA-seq, 44.6 GB for ATAC-seq, and 45.3 GB for ChIP-seq. Individual jobs used up to eight CPU cores. Because the validation datasets differed in the number and size of libraries, these values are intended to provide representative computational requirements rather than direct comparisons among assay types.

## 4 Conclusion

We developed HyARQ, a modular and reproducible Snakemake workflow for allele-specific quantification across RNA-seq, MOA-seq, ATAC-seq, and ChIP-seq data from F1 hybrids. HyARQ enables consistent quantification of parental differences across genes and regulatory regions that are often analyzed using separate approaches. Its quality control outputs help assess data quality and guide filtering and count thresholds before downstream allele-specific analyses. Although HyARQ was evaluated using plant hybrids, the workflow can be applied to F1 hybrids derived from inbred parents in other eukaryotic organisms. It can also be applied to diploid individuals when haplotype-resolved genome assemblies are available. Overall, HyARQ provides a standardized approach for integrated allele-specific analyses across multiple regulatory layers.

## Funding

This work was supported by the National Institute of General Medical Sciences of the National Institutes of Health [grant number R35GM151048]; and the United States Department of Agriculture National Institute of Food and Agriculture [Hatch project 7002327] awarded to J.L.G.

## Acknowledgements

We thank the North Carolina State University High Performance Computing Services Core Facility (RRID: SCR_022168) for providing computational resources and support.

Large language models (LLMs) (Claude Opus 4.8; ChatGPT, GPT-5.6 Sol; OpenAI Codex, GPT-5.6 Terra) were used to assist with scripting, troubleshooting, code documentation, and editing manuscript text for grammar, clarity, and flow. All LLM-assisted outputs were reviewed and verified by the authors.

## Conflicts of interest

None declared.

## Data availability

HyARQ, together with documentation and example test data, is openly available at https://github.com/gagelab/hyarq. The HyARQ v1.0.0 software release is archived in Zenodo under DOI https://doi.org/10.5281/zenodo.23349373. The validation data used in this study, including selected real-data validation inputs and outputs and the complete simulated validation dataset, are available in Zenodo under DOI https://doi.org/10.5281/zenodo.23372517.

## Notes

https://doi.org/10.5281/zenodo.23372517

### Competing Interest Statement

The authors have declared no competing interest.

