## Supplementary Note for "HyARQ: a Snakemake workflow for allele-specific read counting and quality control in F1 hybrids"

### 1 Introduction

HyARQ presents a workflow for allele-specific read counting and quality control across four molecular assays (RNA-seq, MOA-seq, ATAC-seq, and ChIP-seq) in F1hybrids. Analysis of allelic differences in F1 hybrids is widely used to study cis-regulatory effects on variation in gene expression (Hartwig *et al.* 2023, Puixeu *et al.* 2023, Tsouris *et al.* 2023, Engelhorn *et al.* 2025, Huang Y *et al.* 2026). However, allele-specific analyses are sensitive to errors arising from multiple sources, including mapping bias and technical variation, requiring careful processing and quality control (Castel *et al.* 2015). With HyARQ, we aim to facilitate reliable generation of allele-specific counts and provide quality control measures to support downstream analyses. We tested HyARQ using simulated data and applied it to real datasets representing each of the four supported molecular assays. Below, we describe the different steps of the workflow and the testing performed to evaluate its performance and reliability.

### 2 Workflow Steps

HyARQ was designed for allele-specific analysis of sequencing data from F1 hybrids across RNA-seq, MOA-seq, ATAC-seq, and ChIP-seq assays. The workflow consists of five main steps: 1) dual parent reference preparation and paired feature validation, 2) read preprocessing, 3) assay-specific alignment and filtering, 4) feature level counting, and 5) QC reporting. Below, we describe each step in detail.

#### 2.1 Set-up and Input

HyARQ is implemented as a Snakemake workflow and is run using Conda environments that contain the software required for the different workflow steps. The environment files and instructions for installing and running HyARQ are provided with the GitHub repository (https://github.com/gagelab/hyarq). Software used by the workflow and the versions used during development and testing are listed in **Supplementary Table 1**. Once the software environment and workflow configuration are set up, HyARQ can be run through Snakemake using the provided configuration and sample metadata files.

HyARQ requires two parental reference genomes in FASTA format together with paired-end FASTQ files for the sequencing libraries to be analyzed. RNA-seq analysis additionally requires gene annotations in GTF format for the parental genomes and a file defining paired allelic genes between the two parents. MOA-seq, ATAC-seq, and ChIP-seq require files defining paired peak regions representing corresponding allelic loci in the two parental genomes. These paired feature files are supplied by the user and define the allelic loci used for parental counting. The procedures used to generate the paired features for the datasets analyzed in this study are described in the corresponding application sections.

Workflow settings are specified in a YAML configuration file containing the parental reference resources and prefixes, assay-specific options, filtering thresholds, computational resources, and output settings. Sequencing libraries are provided in a tab-delimited sample metadata file with the columns sample id, assay, replicate, path to read 1, and path to read 2. Each row represents one sequencing library and links the sample identifier and assay type to the corresponding paired FASTQ files. HyARQ uses the assay field to determine which assay-specific workflow modules are run. Multiple assay types can be included in the same analysis provided that they use the same pair of parental genomes.

**Supplementary Table 1:** Software used in the HyARQ workflow. Version numbers indicate the versions used during development and testing of HyARQ.

| **Software** | **Version** | **Use in HyARQ** |
|---|---|---|
| Snakemake | 9.23.1 | Workflow management and execution |
| snakemake-executor-plugin-cluster-generic | 1.0.9 | Cluster job submission |
| FastQC | 0.12.1 | Quality assessment of raw and processed sequencing reads for all assays |
| fastp | 1.3.3 | Adapter trimming and quality filtering for RNA-seq, ATAC-seq, and ChIP-seq |
| SeqPurge | ngs-bits 2026_06 | Adapter trimming and quality filtering for MOA-seq |
| FLASH2 | 2.2.00 | Merging overlapping MOA-seq read pairs before alignment |
| STAR | 2.7.11b | Alignment of RNA-seq, MOA-seq, ATAC-seq, and ChIP-seq reads to the concatenated parental reference |
| SAMtools | 1.23.1 | BAM processing and indexing for all assays and duplicate removal for ATAC-seq and ChIP-seq |
| featureCounts (Subread) | 2.1.1 | Feature-level counting for all assays; genes are counted from GTF annotations for RNA-seq, and paired peak regions are converted to SAF format for MOA-seq, ATAC-seq, and ChIP-seq |
| MultiQC | 1.34 | Aggregation of sequencing quality, preprocessing, and alignment reports |

| | | |
|---|---|---|
| Python | 3.12.14 | Paired feature processing, parental count table generation, summary statistics, and QC reporting |
| pandas | 2.3.3 | QC summary processing |
| NumPy | 2.5.3 | Numerical operations for QC/reporting |
| Matplotlib | 3.10.9 | QC figure generation |

**2.2 Dual parent reference preparation and paired feature validation**

HyARQ workflow starts by preparing a concatenated reference from the two parental genomes. Chromosome and contig names in each FASTA file are prefixed with the user-specified Parent1 or Parent2 identifier so that sequences from the two genomes remain distinguishable after concatenation. The renamed parental FASTA files are then combined into a single reference genome used for read alignment. For RNA-seq, sequence names in the parental GTF annotation files are updated with the same prefixes before the annotations are combined.

HyARQ then checks the prepared reference files before downstream analysis. Sequence names must be unique in the concatenated reference and contain the expected parental prefixes. When RNA-seq is included, annotation sequence names are checked against the corresponding parental reference genome to ensure that annotation records do not refer to sequences absent from the FASTA file.

Paired feature files are also validated before counting. RNA-seq uses a two-column table containing corresponding Parent1 and Parent2 gene identifiers. MOA-seq, ATAC-seq, and ChIP-seq use paired genomic coordinates defining corresponding peak regions in the two parental genomes. HyARQ validates these files by checking the required file structure, duplicate entries, and genomic coordinates for paired peak files. The paired features themselves are defined by the user and are not generated by HyARQ. Examples of paired gene and peak region files are available in the HyARQ GitHub repository.

**2.3 Read preprocessing**

HyARQ applies assay-specific preprocessing to the input sequencing reads before alignment. Raw reads from all libraries are first assessed with FastQC (v.0.12.1) (Andrews 2010). RNA-seq, ATAC-seq, and ChIP-seq paired-end reads are then processed with fastp (v.1.3.3) (Chen *et al.* 2018) to remove adapter sequences and filter low-quality reads. Preprocessing parameters are specified separately for each assay in the workflow configuration.

MOA-seq reads are processed following the general preprocessing approach described elsewhere (Liang *et al.* 2022, Engelhorn *et al.* 2025). Because MOA-seq produces short DNA fragments, the paired reads largely overlap and are merged before alignment. HyARQ uses SeqPurge (v.2026_06) (Sturm *et al.* 2016) for adapter and quality trimming, followed by FLASH2 (v.2.2.00) (Magoč and

Salzberg 2011) to merge overlapping read pairs into single-end reads. The merged reads are then used for MOA-seq alignment.

After preprocessing, FastQC is run again for all assay types to assess read quality. MultiQC (v.1.34) (Ewels *et al.* 2016) summarizes the available quality control and alignment reports across all libraries in a single HTML report.

#### 2.4 Assay-specific alignment and filtering

HyARQ uses STAR (v.2.7.11b) (Dobin *et al.* 2013) to align reads from all four assay types to the concatenated parental reference. RNA-seq and MOA-seq mapping settings were based on Engelhorn et al. (2025), ATAC-seq settings on Minow et al. (2025), and ChIP-seq settings on Hartwig et al. (2023). RNA-seq reads are aligned in paired-end mode using splice-aware two-pass mapping with --twopassMode Basic, --outFilterIntronMotifs RemoveNoncanonical, --outFilterType BySJout, --outFilterMultimapNmax 1, --outSAMmultNmax 1, and --winAnchorMultimapNmax 100. Merged MOA-seq reads are aligned as single-end reads using --alignIntronMax 1, --outSAMmultNmax 2, --winAnchorMultimapNmax 100, and --outBAMsortingBinsN 5. ATAC-seq and ChIP-seq reads are aligned in paired-end mode. ATAC-seq uses --alignIntronMax 1, --outSAMmultNmax 2, --winAnchorMultimapNmax 100, and --outBAMsortingBinsN 5, while ChIP-seq uses the STAR settings described by Hartwig et al. (2023), including --alignIntronMax 1. For all assays, only uniquely mapped alignments with a mapping quality of 255 are retained. Duplicate alignments are additionally removed from ATAC-seq and ChIP-seq with SAMtools (v.1.23.1) (Danecek *et al.* 2021) before counting. The final filtered alignments are stored as coordinate-sorted and indexed BAM files.

#### 2.5 Assay-specific parental assignment and feature level counting

HyARQ uses featureCounts (v.2.1.1) (Liao *et al.* 2014) to count parental reads at predefined paired features for all four assay types. For RNA-seq, gene-level counts are generated from the filtered alignments using the concatenated parental GTF annotation. Paired-end fragments are counted according to the strandedness specified in the workflow configuration, and Parent1 and Parent2 gene counts are matched using the supplied gene pair file. For MOA-seq, ATAC-seq, and ChIP-seq, the paired peak regions are converted to SAF format and counted with featureCounts. A minimum overlap of 1 bp between an alignment and a peak region is required for assignment. MOA-seq is counted from merged single-end alignments, whereas ATAC-seq and ChIP-seq are counted as paired-end fragments. Parent1 and Parent2 peak counts are matched according to the supplied peak pair files. For each gene or peak pair, HyARQ reports the Parent1 count, Parent2 count, and total count, and counts from individual libraries are combined into assay-level count matrices.

#### 2.6 QC reporting

HyARQ generates quality control reports at both the sequencing and paired-feature levels. FastQC, fastp, and STAR results are collected across libraries with MultiQC to provide a summary of read quality, preprocessing, and alignment. The reports allow users to examine individual libraries as well as general patterns across an experiment.

For each assay, HyARQ also generates QC summaries from the paired feature count tables. These include the number of paired features with detectable counts, total read or fragment counts assigned to paired features, the proportion of counts assigned to each parent, and the number of paired features retained at selected count thresholds. Summary statistics are provided for individual libraries and across the experiment. HyARQ produces QC tables together with figures in PDF format showing the main count and parental balance metrics. These outputs are intended to allow users to identify low-count libraries, unusual parental count distributions, and changes in the number of informative paired features before carrying out downstream allele-specific analyses. The main count and quality control outputs generated by HyARQ are described in **Supplementary Table 2**.

**Supplementary Table 2:** Description of the main count and quality control outputs generated by HyARQ.

| **Output/metric** | **Description** |
|---|---|
| Sample ID | Identifier for the sequencing library provided in the sample metadata file. |
| Assay | Assay type assigned to the library: RNA-seq, MOA-seq, ATAC-seq, or ChIP-seq. |
| Replicate | Replicate identifier provided in the sample metadata file. |
| Paired features | Number of paired genes or peak regions provided for the assay. |
| Parent1 count | Number of reads or fragments assigned to the Parent1 member of a paired feature. |
| Parent2 count | Number of reads or fragments assigned to the Parent2 member of a paired feature. |
| Total count | Sum of the Parent1 and Parent2 counts for a paired feature. |
| Features with counts | Number of paired features with a total count greater than zero. |
| Total assigned count | Sum of Parent1 and Parent2 counts across all paired features in a library. |
| Parent1 fraction | Proportion of the total assigned count originating from Parent1, calculated as the sum of Parent1 counts across all paired features divided by the total assigned count. |
| Feature retention | Number of paired features retained at the selected minimum total count thresholds. |

| | |
|---|---|
| Paired feature count table | Tab-delimited file containing the Parent1 count, Parent2 count, and total count for each paired gene or peak region in an individual library. |
| Assay-wide count matrix | Combined feature-level counts across all libraries belonging to the same assay. |
| QC summary | Tab-delimited summary of feature detection, count depth, parental count proportions, and other library- and experiment-level QC statistics. |
| QC retention summary | Tab-delimited summary of the number of paired features retained at the selected count thresholds. |
| QC report | PDF report summarizing paired-feature detection, count depth, parental count proportions, and feature retention across samples and the experiment. |
| MultiQC report | HTML report summarizing available read quality, preprocessing, and alignment metrics across libraries. |

### 3 Testing with simulated data

#### 3.1 Simulation methods

To evaluate HyARQ using data with known parental origin and allelic proportions, we generated synthetic datasets for RNA-seq, MOA-seq, ATAC-seq, and ChIP-seq. Two synthetic parental genomes, designated P1 and P2, were constructed with two chromosomes of 100 kb each. P2 was derived from P1 by introducing sequence differences within the simulated features and small insertions and deletions outside the tested features, resulting in corresponding parental loci with sequence and coordinate differences. Four paired features were defined for each assay. RNA-seq features consisted of 1.5-kb genes containing two 500-bp exons separated by a 500-bp intron with canonical GT-AG splice sites in both parents. Parent-specific SNPs were introduced within the exons. ATAC-seq, MOA-seq, and ChIP-seq features were represented by paired regions of 500 bp, 80 bp, and 400 bp, respectively. Parent-specific SNPs were introduced within these regions, including one SNP per MOA-seq peak. Paired gene and peak files were generated in the same formats required as input by HyARQ.

Four parental fractions were simulated within each assay, with Parent1 contributing 0.50, 0.80, 0.20, and 0.50 of fragments for features 1–4, respectively. For each paired feature, 1,000 biological fragments were simulated per replicate, giving 4,000 fragments per library. Two independent replicates were generated for each assay. Reads originating from Parent1 and Parent2 were simulated separately according to the specified parental fraction and subsequently pooled. Read identifiers retained the true parental origin, feature, and replicate to permit evaluation after processing. Fixed random seeds were used to make the simulations reproducible.

RNA-seq transcript sequences were extracted from the parental genome annotations using gffread (v.0.12.9) (Pertea and Pertea 2020), and paired-end reads were generated with ART (v.2016.06.05) (Huang W *et al.* 2012) using the Illumina HiSeq 2500 error model, 150-bp reads, a mean fragment length of 300 bp, and a fragment-length standard deviation of 40 bp. For ATAC-seq, sequences from the predefined paired peak regions were extracted using BEDTools (v.2.31.1) (Quinlan and Hall 2010) and simulated as 150-bp paired-end reads with a mean fragment length of 200 bp and standard deviation of 30 bp. MOA-seq reads were generated from 80-bp paired peak regions as 50-bp paired-end reads with a mean fragment length of 65 bp and standard deviation of 8 bp, producing overlapping mate pairs that could undergo the SeqPurge and FLASH2 preprocessing implemented by HyARQ. ChIP-seq reads were generated from 400-bp paired peak regions as 150-bp paired-end reads with a mean fragment length of 250 bp and standard deviation of 40 bp. No artificial PCR duplicates were added to the ATAC-seq or ChIP- seq simulations. These simulations were designed to test parental read assignment and paired-feature counting rather than reproduce all properties of the individual assays.

The simulated libraries were processed through the complete HyARQ workflow. For each feature and replicate, we calculated parental retention, cross-parent misassignment, final fragment recovery, the observed P1 fraction from the final HyARQ count table, and the absolute difference between observed and expected P1 fractions. ATAC-seq and ChIP-seq parental assignment was evaluated before duplicate removal so that mapping behavior could be distinguished from duplicate filtering, whereas final recovery was calculated from the post-filtering HyARQ feature counts.

#### 3.2 Simulation results

HyARQ recovered the imposed parental proportions for all four assay types. Detailed feature-level results for all assays and replicates are provided in **Supplementary Table 3**. Across the eight simulated feature-replicate combinations per assay, the mean absolute error between expected and observed Parent1 fractions was 0.0020 for RNA-seq, 0.0047 for MOA-seq, 0.0017 for ATAC-seq, and 0.0051 for ChIP-seq. The largest error among all 32 feature-replicate combinations was 0.0231, observed for a MOA-seq feature with an expected Parent1 fraction of 0.20 and an observed Parent1 fraction of 0.2231. Cross-parent assignment was minimal. Among retained fragments, the overall cross-parent misassignment rates were 0% for RNA-seq, 0.113% for MOA-seq, 0.013% for ATAC-seq, and 0.029% for ChIP-seq.

Final fragment recovery differed among assays. Mean recovery was 96.6% for RNA-seq, 77.4% for MOA-seq, 93.8% for ATAC-seq, and 84.9% for ChIP-seq. Recovery also varied among features within an assay. The largest variation occurred for MOA-seq, where the feature with an expected Parent1 fraction of 0.20 had recoveries of 36.3% and 36.2% in the two replicates, while the remaining MOA-seq features ranged from 79.7% to 100%. ATAC-seq recovery ranged from 91.2% to 99.4%. Despite these differences in fragment retention, the expected parental fractions remained closely preserved. For example, the two replicates of the lower-recovery MOA-seq feature produced observed Parent1 fractions of 0.2231 and 0.2017.

Parent1 and Parent2 retention rates were generally similar, and differences varied in direction among features and replicates, indicating no consistent bias toward either parental genome. Cross-parent assignments were also rare at the feature level. No cross-parent assignments were observed for RNA-seq, and for MOA-seq, ATAC-seq, and ChIP-seq, no more than two fragments were assigned to the opposite parent in any feature-replicate combination (**Supplementary Table 3**). These results show that HyARQ preserved the expected parental fractions with minimal cross-parent assignment across all four assay types, even when feature-specific fragment recovery was lower.

**Supplementary Table 3:** Feature-level evaluation of HyARQ using simulated data. Results are shown for four paired features in two independent replicates for RNA-seq, MOA-seq, ATAC-seq, and ChIP-seq. The expected Parent1 fraction indicates the parental proportion specified during simulation, and the observed Parent1 fraction was calculated from the final HyARQ feature counts. Absolute fraction error is the absolute difference between expected and observed Parent1 fractions. Recovery percentage represents the proportion of simulated fragments retained in the final feature counts, while Parent1 and Parent2 retention percentages indicate the proportion of fragments originating from each parent that were retained. Parent1 to Parent2 and Parent2 to Parent1 report the number of fragments assigned to the opposite parental genome. Each feature was simulated with 1,000 fragments per replicate.

| Assay | Replicate | Feature index | Expected parent1 fraction | Observed parent1 fraction | Absolute fraction error | Recovery (%) | Retention parent1 (%) | Retention parent2 (%) | Parent1 to parent2 | Parent2 to parent1 |
|---|---|---|---|---|---|---|---|---|---|---|
| RNA | 1 | 1 | 0.5 | 0.5005 | 0.0005 | 98.9 | 99 | 98.8 | 0 | 0 |
| RNA | 1 | 2 | 0.8 | 0.7956 | 0.0044 | 91.5 | 91 | 93.5 | 0 | 0 |
| RNA | 1 | 3 | 0.2 | 0.2004 | 0.0004 | 96.8 | 97 | 96.75 | 0 | 0 |
| RNA | 1 | 4 | 0.5 | 0.4995 | 0.0005 | 99.7 | 99.6 | 99.8 | 0 | 0 |
| RNA | 2 | 1 | 0.5 | 0.4995 | 0.0005 | 98.5 | 98.4 | 98.6 | 0 | 0 |
| RNA | 2 | 2 | 0.8 | 0.8022 | 0.0022 | 92.5 | 92.75 | 91.5 | 0 | 0 |
| RNA | 2 | 3 | 0.2 | 0.2067 | 0.0067 | 95.8 | 99 | 95 | 0 | 0 |
| RNA | 2 | 4 | 0.5 | 0.499 | 0.001 | 99.4 | 99.2 | 99.6 | 0 | 0 |
| MOA | 1 | 1 | 0.5 | 0.5 | 0 | 100 | 100 | 100 | 0 | 0 |
| MOA | 1 | 2 | 0.8 | 0.8004 | 0.0004 | 92.2 | 92.25 | 92 | 0 | 0 |
| MOA | 1 | 3 | 0.2 | 0.2231 | 0.0231 | 36.3 | 40.5 | 35.25 | 0 | 0 |
| MOA | 1 | 4 | 0.5 | 0.5019 | 0.0019 | 79.7 | 80.4 | 79 | 2 | 0 |
| MOA | 2 | 1 | 0.5 | 0.501 | 0.001 | 100 | 100 | 100 | 0 | 1 |
| MOA | 2 | 2 | 0.8 | 0.8002 | 0.0002 | 92.1 | 92.25 | 91.5 | 1 | 0 |
| MOA | 2 | 3 | 0.2 | 0.2017 | 0.0017 | 36.2 | 36 | 36.25 | 0 | 1 |
| MOA | 2 | 4 | 0.5 | 0.4909 | 0.0091 | 82.3 | 80.4 | 84.2 | 0 | 2 |
| ATAC | 1 | 1 | 0.5 | 0.5 | 0 | 99.4 | 100 | 100 | 0 | 0 |

| | | | | | | | | | | |
|---|---|---|---|---|---|---|---|---|---|---|
| ATAC | 1 | 2 | 0.8 | 0.7996 | 0.0004 | 92.3 | 93.75 | 93.5 | 0 | 0 |
| ATAC | 1 | 3 | 0.2 | 0.2052 | 0.0052 | 92.1 | 95 | 92.5 | 0 | 0 |
| ATAC | 1 | 4 | 0.5 | 0.4973 | 0.0027 | 92.3 | 92.6 | 93.6 | 0 | 0 |
| ATAC | 2 | 1 | 0.5 | 0.5 | 0 | 99.2 | 100 | 100 | 0 | 0 |
| ATAC | 2 | 2 | 0.8 | 0.7963 | 0.0037 | 92.3 | 93.12 | 94.5 | 0 | 0 |
| ATAC | 2 | 3 | 0.2 | 0.1998 | 0.0002 | 91.6 | 92 | 93.12 | 0 | 1 |
| ATAC | 2 | 4 | 0.5 | 0.4989 | 0.0011 | 91.2 | 92.6 | 92.4 | 0 | 0 |
| ChIP | 1 | 1 | 0.5 | 0.5015 | 0.0015 | 96.9 | 97.8 | 97.6 | 0 | 0 |
| ChIP | 1 | 2 | 0.8 | 0.7915 | 0.0085 | 51.8 | 52.25 | 54 | 0 | 0 |
| ChIP | 1 | 3 | 0.2 | 0.2027 | 0.0027 | 95.2 | 96.5 | 96.12 | 0 | 0 |
| ChIP | 1 | 4 | 0.5 | 0.4995 | 0.0005 | 95.1 | 96 | 96.6 | 0 | 0 |
| ChIP | 2 | 1 | 0.5 | 0.5015 | 0.0015 | 97.9 | 98.8 | 98.8 | 0 | 0 |
| ChIP | 2 | 2 | 0.8 | 0.8154 | 0.0154 | 53.1 | 55.25 | 49 | 0 | 0 |
| ChIP | 2 | 3 | 0.2 | 0.2076 | 0.0076 | 94.4 | 99 | 95.62 | 1 | 0 |
| ChIP | 2 | 4 | 0.5 | 0.4974 | 0.0026 | 94.9 | 95.2 | 96.2 | 0 | 1 |

### 4 Testing the pipeline with real data

We applied HyARQ to real RNA-seq, MOA-seq, ATAC-seq, and ChIP-seq datasets to evaluate the workflow across the four supported assay types. The datasets included B73×Mo17 hybrids for RNA-seq, MOA-seq, and ChIP-seq, and B73×CML103 synthetic hybrids for ATAC-seq. For each assay, HyARQ was provided with the required input files, and the workflow successfully completed all processing steps and generated parental count matrices and assay specific QC reports.

For the real datasets, paired features were defined using conservative strategies intended to identify high-confidence corresponding loci between the parental genomes and therefore favor specificity over maximal feature retention. Alternative approaches can define paired regulatory regions using broader syntenic intervals, including loci where a feature is detected in only one parent, and may therefore retain parent-specific regulatory regions that are excluded when a feature is required in both parents. HyARQ does not prescribe a particular method for constructing paired features and can use paired regions generated by different comparative genomic strategies, provided that corresponding coordinates are supplied for both parental genomes.

#### 4.1 Application to RNA-seq data

We analyzed six B73×Mo17 hybrid RNA-seq libraries from Deb et al. (2026) (unpublished data; accession number will be added when available). The B73 v5 and Mo17 CAU v2 genomes, together with their corresponding annotations, were used as the parental references. Gene pairs between the two genomes were identified using the MaizeGDB pan-gene table. We retained one-to-one B73–Mo17 gene relationships, resulting in 24,611 allelic gene pairs.

Across all six libraries, 15,624 gene pairs (63.5%) had at least one parental count, and 11,884 gene pairs had nonzero counts for both parental genes (**Figure S1; Supplementary Table 4**). The number of gene pairs with at least one count ranged from 10,707 to 12,583 among individual libraries. The overall proportion assigned to B73 was 0.493. This proportion was similar among individual libraries, ranging from 0.491 to 0.497. The median B73 proportion across gene pairs was also similar among libraries, ranging from 0.517 to 0.521. These results show consistent parental assignment across the genome among the six RNA-seq libraries.

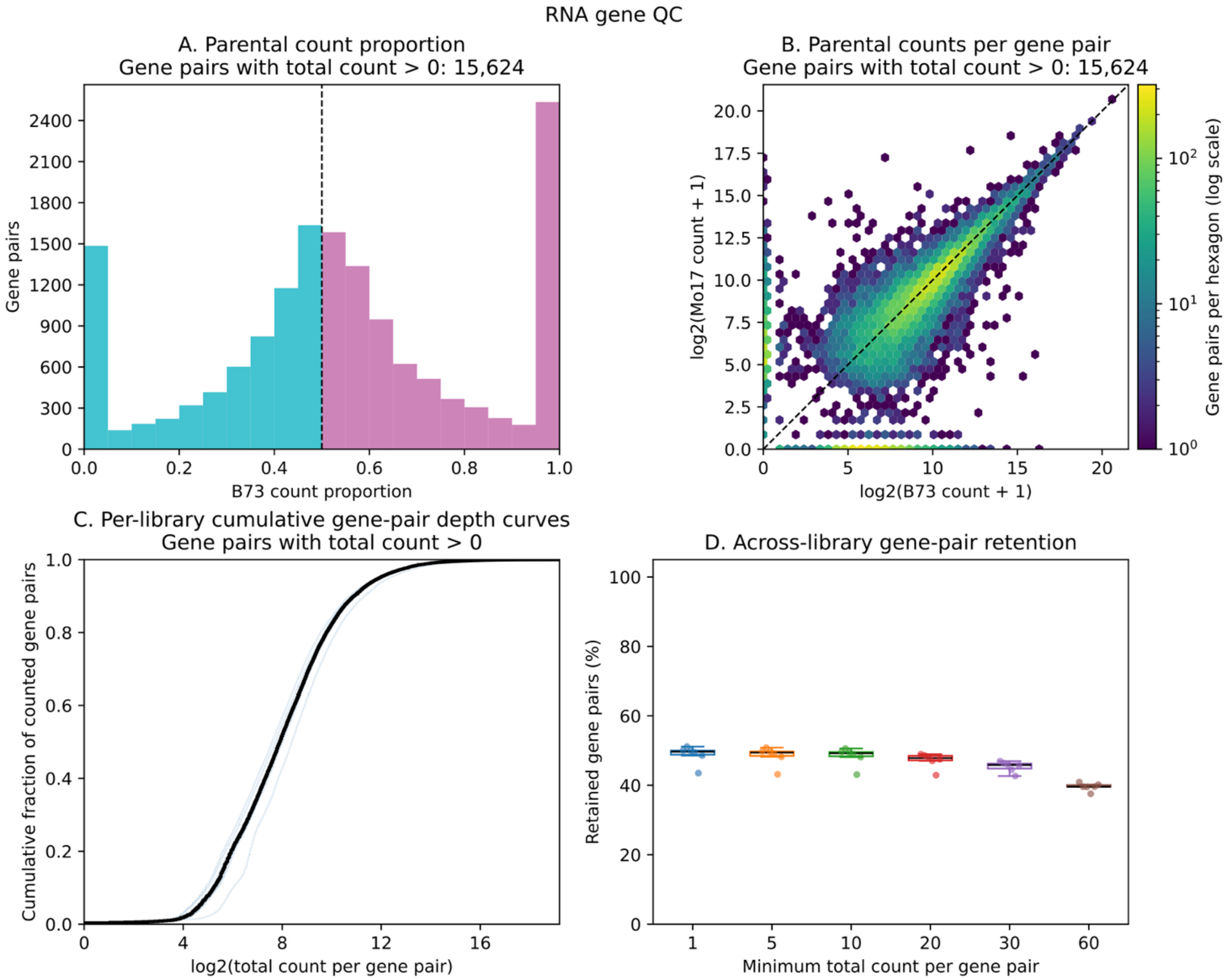


**Figure S1. RNA-seq gene-pair quality control report generated by HyARQ.** Quality control summary for six B73×Mo17 hybrid RNA-seq libraries processed with HyARQ. Across the experiment, 15,624 of the 24,611 supplied gene pairs had a total count > 0. (A) Distribution of the B73 count proportion across gene pairs with a total count > 0. The dashed line indicates equal parental contribution (B73 proportion = 0.5). (B) Relationship between B73 and Mo17 counts for the same gene pairs, shown on a log2(count + 1) scale. The dashed diagonal indicates equal counts between the two parental genes, and hexagon colour represents the number of gene pairs in each bin. (C) Cumulative distributions of gene-pair count depth for the six individual libraries, with total counts shown on a log2 scale. The thick black line represents the median cumulative fraction across libraries at each count depth; the library-specific light-blue curves cluster closely around this median. (D) Percentage of gene pairs retained in each library at minimum total count thresholds of 1, 5, 10, 20, 30, and 60.

**Supplementary Table 4:** RNA-seq gene-pair quality control summary generated by HyARQ. Summary statistics are shown for six B73×Mo17 hybrid RNA-seq libraries and for the combined dataset. Metrics include the total number of supplied gene pairs, the number of gene pairs with counts from either or both parents, total parental counts, overall and median Parent1 count proportions, and the number of gene pairs retained at minimum total count thresholds of 1, 5, 10, 20, 30, and 60. Parent1 corresponds to B73 and Parent2 to Mo17. The all samples column summarizes counts across all six libraries. The HyARQ output table is transposed here for readability.

| | **A10** | **A12** | **A9** | **B11** | **B12** | **C11** | **All samples** |
|---|---|---|---|---|---|---|---|
| **Total gene pairs** | 24611 | 24611 | 24611 | 24611 | 24611 | 24611 | 24611 |
| **Gene pairs with any count** | 12315 | 12583 | 12294 | 12175 | 10707 | 11952 | 15624 |
| **Gene pairs with parent1 count only** | 2261 | 2204 | 2192 | 2207 | 2273 | 2230 | 2381 |
| **Gene pairs with parent2 count only** | 1683 | 1669 | 1536 | 1647 | 1734 | 1688 | 1359 |
| **Gene pairs with both parent counts nonzero** | 8371 | 8710 | 8566 | 8321 | 6700 | 8034 | 11884 |
| **Total parental count** | 14316015 | 15298249 | 14095264 | 15018961 | 18509878 | 14665311 | 91903678 |
| **Overall Parent1 proportion** | 0.494766 | 0.490872 | 0.494719 | 0.49211 | 0.492198 | 0.496555 | 0.493445 |
| **Median gene-pair Parent1 proportion** | 0.518887 | 0.517442 | 0.519371 | 0.518729 | 0.520506 | 0.519675 | 0.524466 |
| **Retained ≥ 1** | 12315 | 12583 | 12294 | 12175 | 10707 | 11952 | 15624 |
| **Retained ≥ 5** | 12248 | 12515 | 12226 | 12102 | 10623 | 11868 | 15522 |
| **Retained ≥ 10** | 12212 | 12452 | 12184 | 12056 | 10601 | 11844 | 15473 |
| **Retained ≥ 20** | 12047 | 11962 | 11857 | 11578 | 10562 | 11689 | 15254 |
| **Retained ≥ 30** | 11556 | 11412 | 11324 | 10952 | 10499 | 11247 | 14899 |
| **Retained ≥ 60** | 10074 | 9742 | 9727 | 9239 | 9747 | 9890 | 13851 |

#### 4.2 Application to MOA-seq data

To evaluate HyARQ with MOA-seq data, we used three publicly available B73×Mo17 MOA-seq paired-end libraries from Engelhorn et al. (2025): SRR28762725, SRR28762726, and SRR28762727. The B73 NAM v5 and Mo17 CAU v1 genomes were used as the parental references, consistent with the genomes used in the published study. Because HyARQ requires predefined paired peak regions between parental genomes, we generated a set of one-to-one B73–Mo17 peak pairs from the published MOA peaks.

The published MOA dataset contained 327,029 peaks; after assignment to the parental genomes and restriction to the ten primary chromosomes, 166,948 B73 peaks and 157,720 Mo17 peaks were considered for pairing. Whole-genome alignments between B73 and Mo17 were generated using the proali function of AnchorWave (v.1.2.6) (Song *et al.* 2022), with anchor regions identified from the B73 gene annotation using minimap2 (v.2.31) (Li 2018). The resulting MAF alignment was converted to chain format using wgatools (v.1.1.0) (Wei *et al.* 2025) and used with CrossMap (v.0.7.4) (Zhao *et al.* 2014) for bidirectional liftover between the two genomes. We used a conservative pairing strategy designed to retain only high-confidence one-to-one correspondences. B73 peaks were lifted to the Mo17 genome, and BEDTools (v.2.31.1) (Quinlan and Hall 2010) was used to identify overlaps with Mo17 peaks. Candidate pairs were retained only when the lifted region mapped as a single continuous interval and overlapped a single peak in the target genome. The procedure was then repeated in the reverse direction, and only reciprocal pairs that mapped back to the corresponding original region were retained. This approach prioritizes unambiguous correspondence between parental regulatory regions rather than maximizing the number of retained peaks and produced 8,242 one-to-one B73–Mo17 peak pairs.

The resulting 8,242 one-to-one B73–Mo17 peak pairs were used as input for HyARQ. All 8,242 peak pairs received at least one count in each of the three libraries (**Figure S2; Supplementary Table 5**). The overall B73 proportion was 0.500. Individual libraries showed similarly balanced parental proportions of 0.499, 0.502, and 0.499. The median B73 proportion across paired peaks was 0.500 for each library. Overall, HyARQ recovered parental counts across the complete set of paired MOA-seq peaks, with similar representation of the two parental genomes.

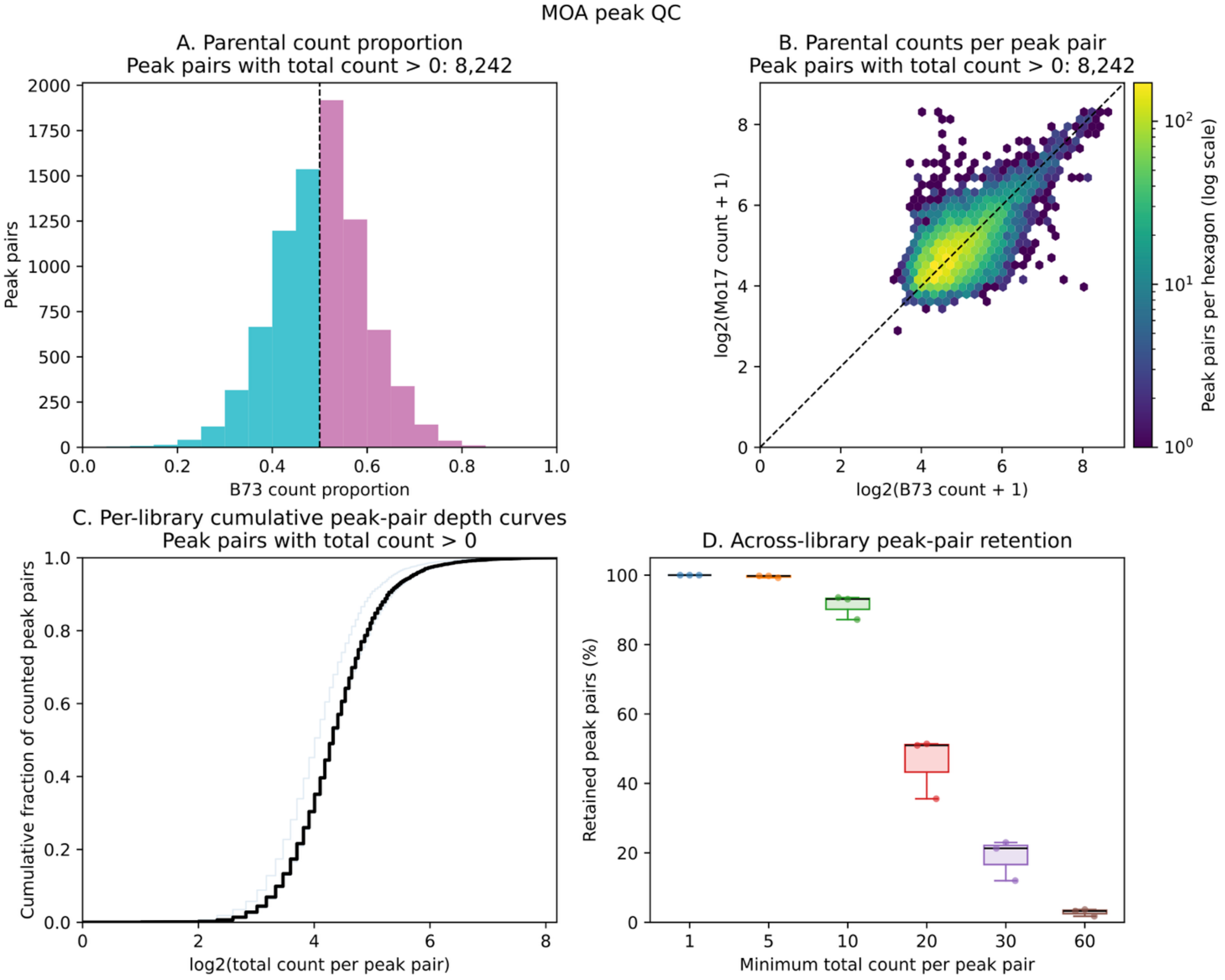


**Figure S2. MOA-seq peak-pair quality control report generated by HyARQ.** Quality control summary for three B73×Mo17 hybrid MOA-seq libraries processed with HyARQ. Across the experiment, all 8,242 supplied peak pairs had a total count > 0. (A) Distribution of the B73 count proportion across counted peak pairs. The dashed line indicates equal parental contribution (B73 proportion = 0.5). (B) Relationship between B73 and Mo17 counts for the same peak pairs, shown on a log2(count + 1) scale. The dashed diagonal indicates equal counts between the two parental peak regions, and hexagon colour represents the number of peak pairs in each bin. (C) Per-library cumulative peak-pair depth curves showing the distribution of total counts across counted peak pairs, with individual libraries shown in light blue and the median cumulative distribution across libraries shown in black. (D) Percentage of peak pairs retained in individual libraries at minimum total count thresholds of 1, 5, 10, 20, 30, and 60.

**Supplementary Table 5:** MOA-seq peak-pair quality control summary generated by HyARQ. Summary statistics are shown for three B73×Mo17 hybrid MOA-seq libraries and for the combined dataset. Metrics include the total number of supplied peak pairs, the number of peak pairs with counts from either or both parents, total parental counts, overall and median Parent1 count proportions, and the number of peak pairs retained at minimum total count thresholds of 1, 5, 10, 20, 30, and 60. Parent1 corresponds to B73 and Parent2 to Mo17. The all samples column summarizes counts across all three libraries. The HyARQ output table is transposed here for readability.

| | **SRR28762725** | **SRR28762726** | **SRR28762727** | **All samples** |
|---|---|---|---|---|
| **Total peak pairs** | 8242 | 8242 | 8242 | 8242 |
| **Peak pairs with any count** | 8242 | 8242 | 8242 | 8242 |
| **Peak pairs with parent1 count only** | 17 | 21 | 37 | 0 |
| **Peak pairs with parent2 count only** | 17 | 19 | 39 | 0 |
| **Peak pairs with both parent counts nonzero** | 8208 | 8202 | 8166 | 8242 |
| **Total parental count** | 196791 | 201300 | 160088 | 558179 |
| **Overall Parent1 proportion** | 0.498798 | 0.50152 | 0.499044 | 0.49985 |
| **Median peak-pair Parent1 proportion** | 0.5 | 0.5 | 0.5 | 0.5 |
| **Retained ≥ 1** | 8242 | 8242 | 8242 | 8242 |
| **Retained ≥ 5** | 8225 | 8223 | 8179 | 8242 |
| **Retained ≥ 10** | 7709 | 7672 | 7187 | 8242 |
| **Retained ≥ 20** | 4200 | 4235 | 2933 | 8241 |
| **Retained ≥ 30** | 1756 | 1894 | 988 | 8124 |
| **Retained ≥ 60** | 267 | 302 | 144 | 3558 |

#### 4.3 Application to ATAC-seq data

To our knowledge, no publicly available plant hybrid ATAC-seq dataset has both parental genome assemblies available. We therefore used publicly available B73 and CML103 inbred ATAC-seq data from Hufford et al. (2021) to generate four synthetic B73×CML103 libraries. The B73 libraries were SRR14327895, SRR14327896, SRR14327897, and SRR14327898, and the CML103 libraries were SRR14327903, SRR14327904, SRR14327905, and SRR14327906. Each synthetic library was created by combining equal numbers of paired-end reads randomly sampled from matched B73 and CML103 libraries: SRR14327895 with SRR14327903 (B73×CML103 rep1), SRR14327896 with SRR14327904 (B73×CML103 rep2), SRR14327897 with SRR14327905 (B73×CML103 rep3), and SRR14327898 with SRR14327906 (B73×CML103 rep4).

The B73 NAM v5 and CML103 NAM v1 genome assemblies were used as the parental references. Published ATAC-seq peak files in the native coordinates of each genome were obtained from Hufford et al. (2021), containing 50,486 B73 and 34,326 CML103 accessible chromatin regions (ACRs). After restriction to the ten primary chromosomes, 50,481 B73 and 34,261 CML103 ACRs were considered for pairing. Paired ACRs were identified using a modified version of the reciprocal ACR-alignment approach described by Minow et al. (2025). ACR sequences were extracted with BEDTools and aligned to the opposite parental genome using minimap2. Candidate alignments were filtered using one-to-one pangene relationships from the maize pan-genome annotation, retaining alignments in which the query ACR and target genomic interval shared a corresponding pangene. Alignment blocks were then intersected with the ACR set from the target parent using BEDTools, and candidate pairs were retained when the summed overlap covered at least 45% of the target ACR. The procedure was performed in both directions, and only strict reciprocal one-to-one pairs were retained, such that each B73 ACR had a single CML103 partner and that CML103 ACR reciprocally had the same B73 ACR as its single partner. This procedure produced 14,302 B73–CML103 ACR pairs, which were used as the paired peak input for HyARQ.

Across the four B73×CML103 libraries analyzed using 14,302 paired ACRs, 12,300 paired regions (86.0%) had at least one parental count, and 11,469 had nonzero counts for both parental ACRs (**Figure S3; Supplementary Table 6**). The number of paired regions with counts in individual libraries ranged from 11,085 to 11,665. The overall B73 proportion was 0.534, with individual library proportions of 0.448, 0.485, 0.576, and 0.587. The corresponding median B73 proportions across paired regions ranged from 0.450 to 0.600. Because the controlled simulation showed no systematic difference in parental assignment, the variation among these synthetic libraries is consistent with differences in source-library coverage and the distribution of reads across paired ACRs rather than a consistent parental bias introduced by HyARQ.

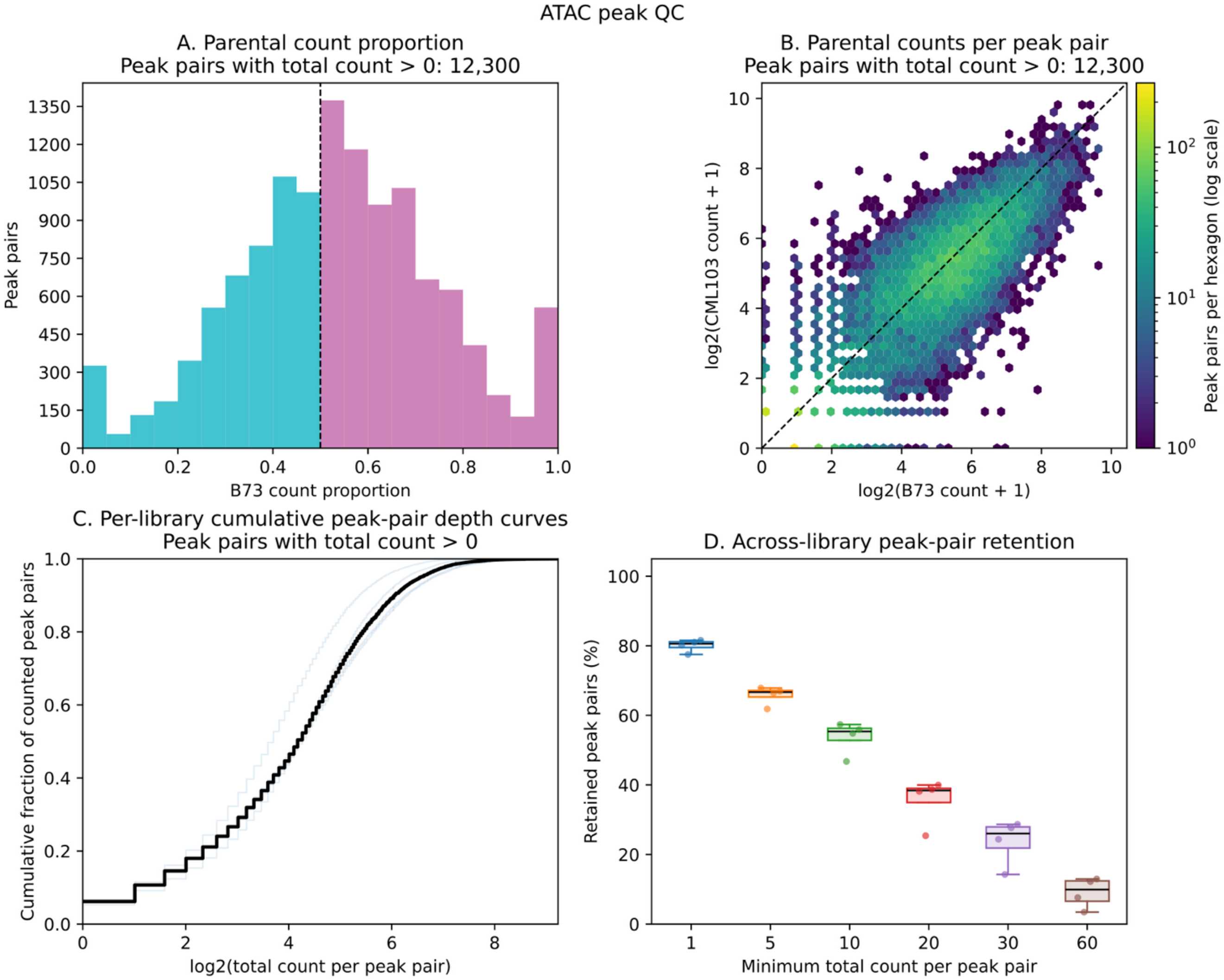


**Figure S3. ATAC-seq peak-pair quality control report generated by HyARQ.** Quality control summary for four synthetic B73×CML103 hybrid ATAC-seq libraries processed with HyARQ. Across the experiment, 12,300 of the 14,302 supplied peak pairs had a total count > 0. (A) Distribution of the B73 count proportion across counted peak pairs, with the dashed line indicating equal parental contribution. (B) Hexbin plot comparing B73 and CML103 counts for individual peak pairs on a log2 scale; the dashed diagonal indicates equal counts in the two parents, and color indicates the number of peak pairs per hexagon. (C) Per-library cumulative peak-pair depth curves showing the distribution of total counts across counted peak pairs, with individual libraries shown in light blue and the median cumulative distribution across libraries shown in black. (D) Across library peak-pair retention at increasing minimum total count thresholds, summarized as the percentage of supplied peak pairs retained in each library.

**Supplementary Table 6:** ATAC-seq peak-pair quality control summary generated by HyARQ. Summary statistics are shown for four synthetic B73×CML103 hybrid ATAC-seq libraries and for the combined dataset. Metrics include the total number of supplied peak pairs, the number of peak pairs with counts from either or both parents, total parental counts, overall and median Parent1 count proportions, and the number of peak pairs retained at minimum total count thresholds of 1, 5, 10, 20, 30, and 60. Parent1 corresponds to B73 and Parent2 to CML103. The all samples column summarizes counts across all four libraries. The HyARQ output table is transposed here for readability.

| | **B73×CML103 rep1** | **B73×CML103 rep2** | **B73×CML103 rep3** | **B73×CML103 rep4** | **All samples** |
|---|---|---|---|---|---|
| **Total peak pairs** | 14302 | 14302 | 14302 | 14302 | 14302 |
| **Peak pairs with any count** | 11465 | 11085 | 11590 | 11665 | 12300 |
| **Peak pairs with parent1 count only** | 610 | 831 | 1102 | 1115 | 532 |
| **Peak pairs with parent2 count only** | 772 | 947 | 511 | 525 | 299 |
| **Peak pairs with both parent counts nonzero** | 10083 | 9307 | 9977 | 10025 | 11469 |
| **Total parental count** | 299048 | 205479 | 371796 | 388790 | 1265113 |
| **Overall Parent1 proportion** | 0.448333 | 0.484575 | 0.576362 | 0.586882 | 0.534423 |
| **Median peak-pair Parent1 proportion** | 0.45 | 0.5 | 0.592413 | 0.6 | 0.531646 |
| **Retained ≥ 1** | 11465 | 11085 | 11590 | 11665 | 12300 |
| **Retained ≥ 5** | 9706 | 8843 | 9501 | 9570 | 11249 |
| **Retained ≥ 10** | 8206 | 6685 | 7842 | 7996 | 10616 |
| **Retained ≥ 20** | 5452 | 3632 | 5534 | 5715 | 9706 |
| **Retained ≥ 30** | 3485 | 2039 | 3957 | 4099 | 8889 |
| **Retained ≥ 60** | 1087 | 490 | 1746 | 1853 | 6570 |

#### 4.4 Application to ChIP-seq data

We applied HyARQ to three publicly available B73×Mo17 hybrid ZmBZR1 ChIP-seq libraries from Hartwig et al. (2023): SRR22541773, SRR22541781, and SRR22541782. The B73 NAM v5 and Mo17 CAU v1 genomes were used as the parental references. Paired B73 and Mo17 ZmBZR1 binding regions were generated from the BZR1 sites reported in the study. The published peak set contained 52,765 parental peak records, including 27,289 B73 and 25,476 Mo17 regions. To identify corresponding binding regions between the two genomes, we used the same reciprocal pairing procedure described for the MOA-seq peaks. This resulted in 1,927 one-to-one B73–Mo17 BZR1 peak pairs, which were used as input to HyARQ.

Across the three libraries, all 1,927 peak pairs received at least one parental count, and nearly all had counts from both parents (**Figure S4; Supplementary Table 7**). Two libraries contained nonzero counts from both parents at all 1,927 peak pairs, while the third contained counts from both parents at 1,925 peak pairs. The overall B73 proportion was 0.493, with individual library proportions ranging from 0.490 to 0.494. The median B73 proportion across peak pairs was approximately 0.500 for all three libraries. Overall, HyARQ showed consistent parental assignment across the three ChIP-seq libraries and recovered counts across nearly all supplied paired BZR1 peaks.

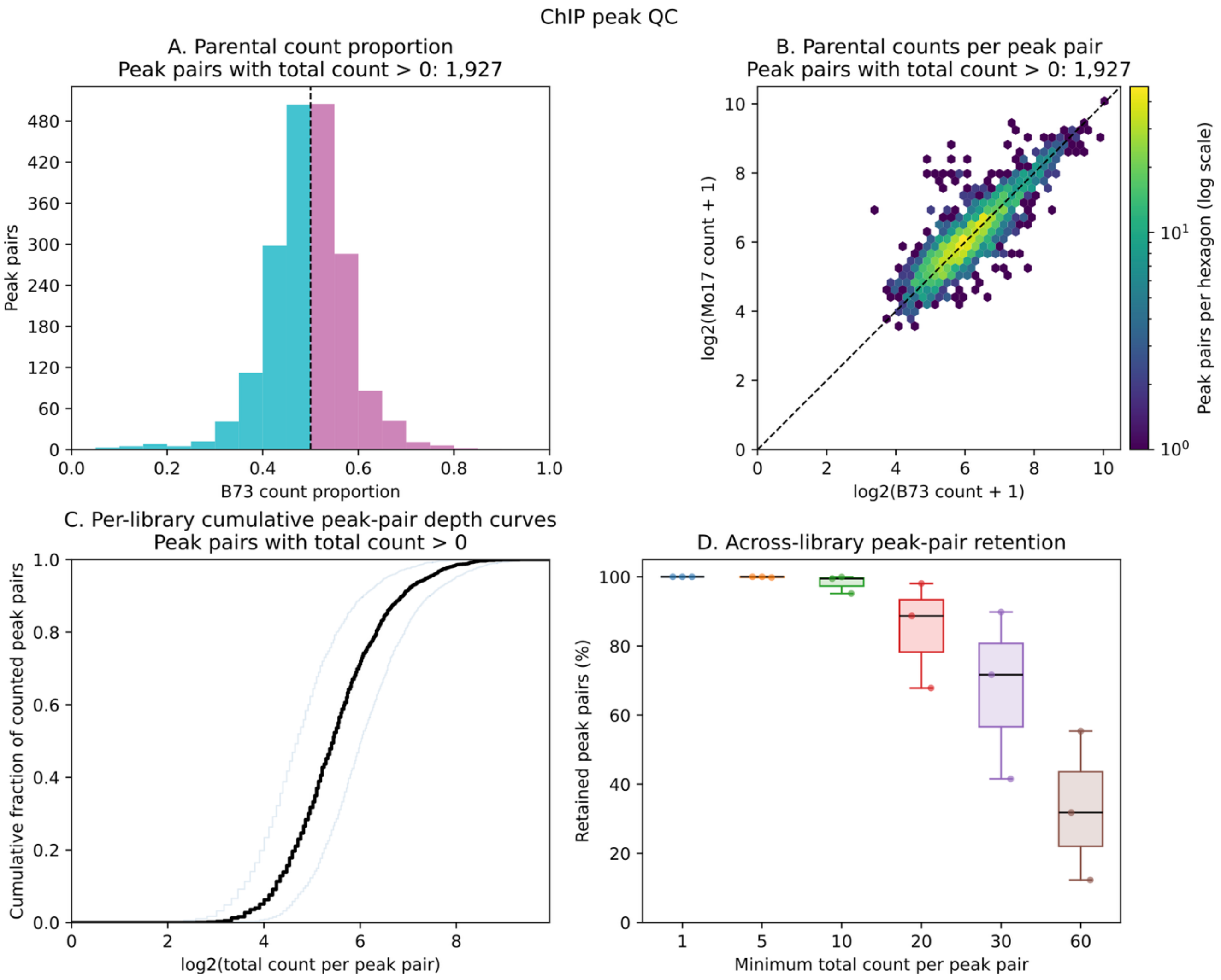


**Figure S4. ChIP-seq peak-pair quality control report generated by HyARQ.** Quality control summary for three B73×Mo17 hybrid ZmBZR1 ChIP-seq libraries processed with HyARQ. Across the experiment, all 1,927 supplied peak pairs had a total count > 0. (A) Distribution of the B73 count proportion across counted peak pairs, with the dashed line indicating equal parental contribution. (B) Hexbin plot comparing B73 and Mo17 counts for individual peak pairs on a log2 scale; the dashed diagonal indicates equal counts in the two parental peak regions, and color indicates the number of peak pairs per hexagon. (C) Per-library cumulative peak pair depth curves showing the distribution of total counts across counted peak pairs, with individual libraries shown in light blue and the median cumulative distribution across libraries shown in black. (D) Across library peak pair retention at increasing minimum total count thresholds, summarized as the percentage of supplied peak pairs retained in each library. Together, these plots summarize parental count balance, count depth, and the effect of minimum count thresholds on the number of peak pairs available for downstream allele-specific analysis.

**Supplementary Table 7:** ChIP-seq peak-pair quality control summary generated by HyARQ. Summary statistics are shown for three B73×Mo17 hybrid ZmBZR1 ChIP-seq libraries and for the combined dataset. Metrics include the total number of supplied peak pairs, the number of peak pairs with counts from either or both parents, total parental counts, overall and median Parent1 count proportions, and the number of peak pairs retained at minimum total-count thresholds of 1, 5, 10, 20, 30, and 60. Parent1 corresponds to B73 and Parent2 to Mo17. The all samples column summarizes counts across all three libraries. The HyARQ output table is transposed here for readability.

| | **SRR22541773** | **SRR22541781** | **SRR22541782** | **All samples** |
|---|---|---|---|---|
| **Total peak pairs** | 1927 | 1927 | 1927 | 1927 |
| **Peak pairs with any count** | 1927 | 1927 | 1927 | 1927 |
| **Peak pairs with parent1 count only** | 0 | 0 | 1 | 0 |
| **Peak pairs with parent2 count only** | 0 | 0 | 1 | 0 |
| **Peak pairs with both parent counts nonzero** | 1927 | 1927 | 1925 | 1927 |
| **Total parental count** | 114722 | 174172 | 67454 | 356348 |
| **Overall Parent1 proportion** | 0.490011 | 0.494442 | 0.493418 | 0.492822 |
| **Median peak-pair Parent1 proportion** | 0.5 | 0.5 | 0.5 | 0.496063 |
| **Retained ≥ 1** | 1927 | 1927 | 1927 | 1927 |
| **Retained ≥ 5** | 1927 | 1927 | 1923 | 1927 |
| **Retained ≥ 10** | 1917 | 1926 | 1834 | 1927 |
| **Retained ≥ 20** | 1709 | 1890 | 1307 | 1927 |
| **Retained ≥ 30** | 1381 | 1731 | 801 | 1925 |
| **Retained ≥ 60** | 613 | 1067 | 237 | 1773 |

Overall, we have shown that HyARQ produces reliable allele-specific count estimates across simulated and published RNA-seq, MOA-seq, ATAC-seq, and ChIP-seq datasets. We provide an open-source Snakemake pipeline, HyARQ, to aid researchers in the analysis of allele-specific molecular data from F1 hybrids and to improve reproducibility and consistency across assays and studies.
